# Oral microbial dysbiosis linked to worsened periodontal condition in rheumatoid arthritis patients

**DOI:** 10.1101/450056

**Authors:** Jôice Dias Corrêa, Gabriel R. Fernandes, Débora Cerqueira Calderaro, Santuza Maria Souza Mendonça, Janine Mayra Silva, Mayra Laino Albiero, Fernando Q Cunha, E Xiao, Gilda Aparecida Ferreira, Antônio Lúcio Teixeira, Chiranjit Mukherjee, Eugene J. Leys, Tarcília Aparecida Silva, Dana T. Graves

## Abstract

Rheumatoid arthritis (RA) is an autoimmune disorder associated with increased periodontal destruction. It is thought that RA increases the risk of periodontal disease; it is not known how it influences the oral microbiota. Our aim was to analyze the impact of RA on subgingival microbiota and its association with periodontal inflammation and RA activity. Forty-two patients with RA were compared to 47 control subjects without RA. Patients were screened for probing depth, clinical attachment level, bleeding on probing and classified as with or without periodontitis. Subgingival plaque was examined by Illumina MiSeq Sequencing of 16S rRNA gene V4 region and inflammatory cytokines were measured in saliva. RA was associated to severe periodontal disease. In addition, the severity of RA, reflected by the number of tender and swollen joints, was significantly correlated with the presence of pathogenic oral bacteria (i.e. *Fusobacterium nucleatum* and *Treponema socransky*). Non-periodontitis RA patients compared to healthy controls had increased microbial diversity and bacterial load, higher levels of pathogenic species (*Prevotella, Selenomonas, Anaeroglobus geminatus, Parvimonas micra, Aggregatibacter actinomycetemcomitans*) and reduction of health-related species (*Streptococcus, Rothia aeria, Kingela oralis*). Genes involved with bacterial virulence (i.e. lipopolysaccharide biosynthesis, peptidases) were more prevalent in the subgingival metagenome of subjects with RA. In addition, the degree of oral inflammation reflected by IL-2, IL-6, TNF-α, IFN-γ salivary levels was increased in non-periodontitis RA patients in comparison with controls. Our findings support the hypothesis that RA triggers dysbiosis of subgingival microbiota, which may contribute to worsening periodontal status.

**Author Summary:** Rheumatoid arthritis (RA) is an autoimmune disease characterized by joints inflammation, swelling, pain and stiffness. Exactly what starts this disease is still unclear. Some recent studies have suggested mucosal surfaces in the body, like those in the gums, could affect the disease process. It has been observed that people with RA have higher risk of periodontitis (a bacterial inflammatory disease of the gums), compared with the general population, and this may be the start of the autoimmune process. Also, periodontitis increases the severity of RA while interventions by treating periodontitis can improve the symptoms of RA. One of the possible mechanisms that link the higher prevalence of periodontitis in RA patients is the dysbiosis of the oral microbiota triggered by the chronic inflammation in RA. Increased levels of molecules of inflammation may affect the oral environment and change the type of bacteria that live there. Here, we examined RA patients and healthy subjects, screening their oral health and inflammatory markers. We collected their saliva and the dental plaque from the space between the teeth and the gum. We found that RA patients exhibited severe periodontitis, increased levels of inflammatory mediators on their saliva and distinct bacterial communities, with higher proportions of bacteria species linked to periodontal disease, even in patients without periodontitis. We also found that the presence of these bacteria species was linked to worse RA conditions. Our study provides new insights to understand the bi-directional mechanisms linking periodontal disease to the development of RA, showing that we need to pay attention to the oral cavity in patients with RA and refer people for dental evaluation. This practice might have a positive impact in the course of RA.

## Introduction

The oral cavity is the second largest microbial niche after the gastrointestinal tract with over 700 bacterial species [1]. In periodontally healthy individuals, microbial populations co-exist in equilibrium with the host. The change in this equilibrium is linked to the pathogenesis of oral diseases such as periodontitis [1]. Oral bacteria, which exist as a biofilm on the tooth surface, can induce inflammation in the adjacent gingiva, leading to osteoclast formation and bone loss which, in severe cases causes tooth loss [2]. Systemic inflammatory diseases may contribute to disrupting the balance between host and oral microbiota [3].

Rheumatoid arthritis (RA) is a systemic autoimmune disease characterized by chronic inflammation and damage to soft and hard articular tissues [5]. An increased incidence of periodontitis has been reported in patients with RA [6]. Furthermore, treatment of periodontitis has been shown to reduce RA activity [7]. A link between periodontal disease and RA involves the production of enzymes capable of modifying proteins to enhance their antigenicity by the addition of malondialdehyde-acetaldehyde, citrullination and carbamylation [8]. Furthermore, RA enhance systemic inflammation which can amplify the local inflammatory response in the periodontium, increasing periodontal destruction [9].

Few studies have described the composition of the oral microbiota in patients with RA [10–13]. Zhang examined dental and salivary microbiome but the periodontal status of RA subjects was not defined [11]. Another study evaluated subgingival microbiota and the periodontal condition of RA subjects, but it did not evaluate the impact of RA and periodontitis independently [10]. Mikuls compared RA to Osteoarthritis patients [12] while Lopez-Oliva analyzed only RA patients without periodontitis [13]. Furthermore, neither of these studies assessed inflammatory parameters in the oral cavity. Our study characterized the subgingival microbiome of RA patients and its association with periodontal status, inflammatory markers and RA scores to establish a link between these parameters.

## Results

### Periodontal destruction and RA outcomes

Of the 42 patients with RA included in the study, 50% had periodontitis compared to 42.6% of the controls (P>0.05). The mean duration of RA was similar for patients without periodontitis (16.18±8.2 years) and for those with periodontitis (12.46±9.7years) (P>0.05). RA activity parameters (number of tender and swollen joints, DAS-28) and medications in use were not different between RA patients with or without periodontitis (Table 1). Of note, the majority of RA subjects (85.7%) with periodontitis were positive for the presence of autoantibodies (ACPA) compared to only 33% in RA patients without periodontitis (P<0.05).

**Table 1.** Demographic and clinical data of patients with RA and healthy control subjects.

|  | Controls |  | RA |  |
| --- | --- | --- | --- | --- |
|  | Non-CP | CP | Non-CP | CP |
| Subjects | 27(57.4%) | 20(42.6%) | 21(50%) | 21(50%) |
| Females % | 36.5 | 48 | 53.48 | 34 |
| Age, years | 42.8(±14.0) | 46.5(±12.3) | 50(±11.1) | 53(±10.40) |
| Current Smokers | 1(3.7%) | 2(7.6%) | 1(4.6%) | 1(4.6%) |
| RA duration, years | n/a | n/a | 16.2(±8.2) | 12.5(±9.7) |
| **Disease active parameters** |  |  |  |  |
| Tender joints | n/a | n/a | 3.2(±4.0) | 3.3(±4.6) |
| Swollen joints | n/a | n/a | 2.5(±0.5) | 2.4(±0.9) |
| DAS28 | n/a | n/a | 3.5(±1.2) | 3.7(±1.5) |
| **Autoantibody status** |  |  |  |  |
| ACPA positive, % | n/a | n/a | 7(33%) | 85.7* |
| **Medications** |  |  |  |  |
| Methotrexate | n/a | n/a | 11(52.4%) | 14(66.7%) |
| Prednisone | n/a | n/a | 9(42.9%) | 14(66.7%) |
| Biological agent | n/a | n/a | 5(23.8%) | 4(19.0%) |
| **Periodontal parameters** |  |  |  |  |
| PD (mm) | 1.9(1.6-2.2) | 3.0(3-3.7)* | 2.9(2-3)<sup>#</sup> | 3.8(3.4-4.5)*<sup>#</sup> |
| CAL (mm) | 2.2(2-3) | 3.0(2.6-3.5) | 3.0(2.9-3.4) | 4.0(3.7-5.3)*<sup>#</sup> |
| BOP (% sites) | 6(1.2-16) | 6.7(2.7-13) | 5(3.8-7.7) | 7(5-14) |
| Missing teeth | 2(0-6) | 4(1-7) | 6(2.5-11)<sup>#</sup> | 6(3-10) |
| Plaque Index | 0.5(0.1-1) | 0.5(0.3-0.8) | 0.5(0.2-0.7) | 0.58(0.23-1.1) |
| Tooth brushing (times/day) | 2.85(±0.93) | 2.62(±0.85) | 2.82(±0.5) | 2.69(±0.6) |

Values were expressed in mean ± SD or median (25% percentile-75% percentile)
CP: Chronic Periodontitis, BOP: bleeding upon probing, PD probing depth, CAL clinical attachment level, DAS28: Disease Activity Score, ACPA anti-citrullinated protein antibody.
\*Statistically different comparing Non-CP x CP within the same group
<sup>#</sup> Statistically different comparing RA x Healthy Control group
One Way ANOVA or Kruskal-Wallis test, p<0.05

The presence of RA was associated with worse periodontal parameters compared with control subjects: probing depth (Controls 3.0 x RA 3.8 mm) and clinical attachment loss (Controls 3.0 x RA 4 mm) (Table 1). These data indicate severe periodontitis in RA patients although self-reported hygiene habits and plaque index did not differ among RA and control subjects.

### RA affects subgingival microbial load, richness and diversity

We investigated the total microbial biomass in RA subjects and found that non-periodontitis RA patients had a significantly ∼1 log higher bacterial burden than did control individuals without periodontitis (Fig 1A). A total of 779 OTUS were found and the impact of RA status on microbial diversity and richness was examined by assessing the number of observed OTUs, Chao1 and Shannon indexes. RA patients had increased microbial diversity compared to controls, both without periodontitis. In subjects with periodontitis, RA was associated with increased diversity assessed by number of OTUs (Fig 1C) and Shannon Index (Fig 1D). Thus, RA was associated with an increased diversity like that one observed in control patients with periodontitis compared to control patients without periodontitis (Fig 1B and 1C).

**Figure 1.**
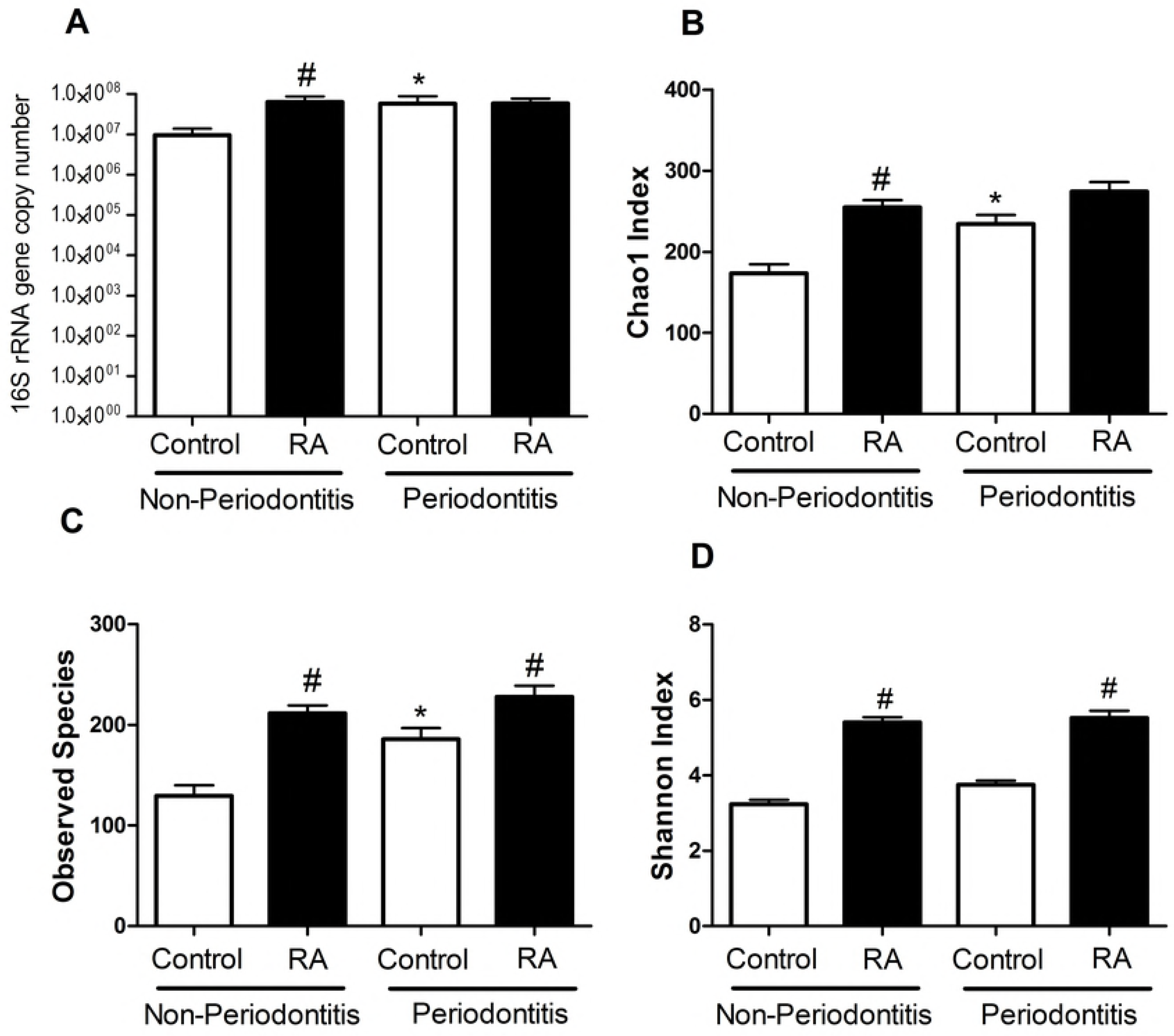
Bacterial load and microbial diversity in subgingival biofilm samples. A) bacterial load, B, C and D) metrics of alpha diversity in Control subjects and RA patients with or without periodontitis. *Statistically different compared to Non-Periodontitis subjects within the same group. #statistically different compared to Controls. p<0.05, Kruskal-Wallis

To analyze whether the subgingival microbial communities in patients with RA were distinct from that of controls, we performed unweighted UniFrac distance analysis (Fig 2). Microbial communities in patients with RA had distinct clusters compared to control patients without the complicating factor of periodontitis (Fig 2A, PERMANOVA, p<0.01). The presence of periodontitis in patients with RA obviated the difference between the RA and control group (P>0.05, Fig 2B). However, RA patients with periodontitis clustered separately from RA patients without periodontitis (P<0.05, Fig 2C).

**Figure 2.**
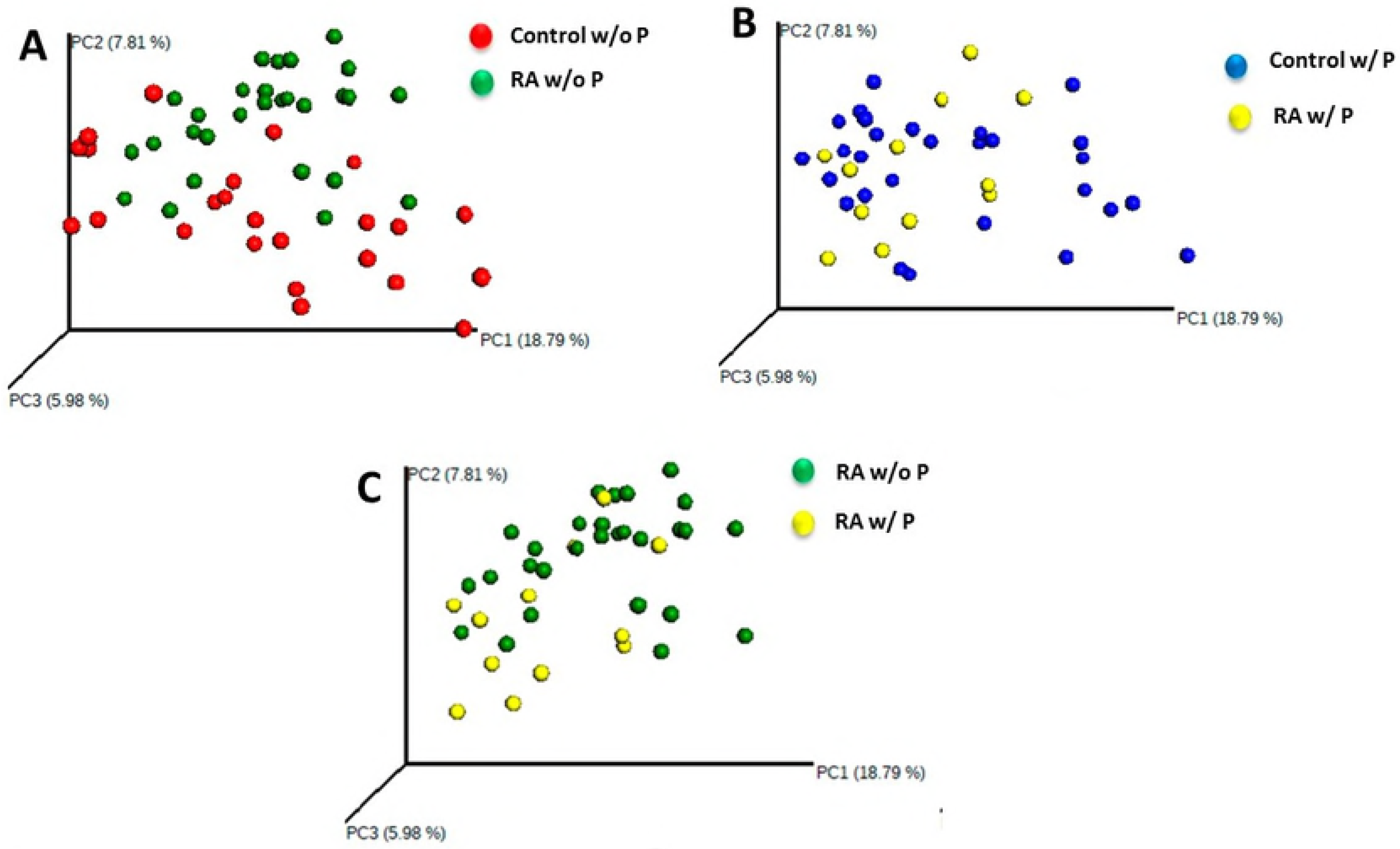
Principal coordinates analysis plot showing the comparison of subgingival microbial community composition. Each point represents a subject. (A) Microbial communities in Control and RA subjects without periodontitis. (w/o P). (B) Microbial communities in RA patients versus Control subjects with periodontitis (w/P). (C) Microbial communities in RA patients without periodontitis versus with periodontitis.

### Subgingival RA and Control group sites harbor distinct bacterial communities

We performed LEFSE (linear discriminant analysis coupled with effect size measurements) for analysis of the relative abundance of microbial taxonomic groups. A number of pathogenic bacteria were significantly elevated in the RA group that are associated with worse periodontal status [1]. RA patients without periodontitis had enrichment in periodontitis-associated bacteria such as *Prevotella* species (*P. melaninogenica, P. denticola, P. histicola, P. nigrescens, P. oulorum,* and *P. maculosa*) and other pathogenic species (*Selenomonas noxia, S. sputigena and Anaeroglobus geminatus*). In addition, RA subjects presented a reduction of health-associated species (*Streptococcus, Rothia aeria, Kingella oralis, Haemophilus, Actinomyces*) (Fig 3A). In the same way, pathogenic species such as *Prevotella, Aggregaticbacter actinomycetemcomitans* and *Parvimonas micra* were significantly increased in RA patients with periodontitis compared to control subjects with periodontitis (Fig 3B).

**Figure 3.**
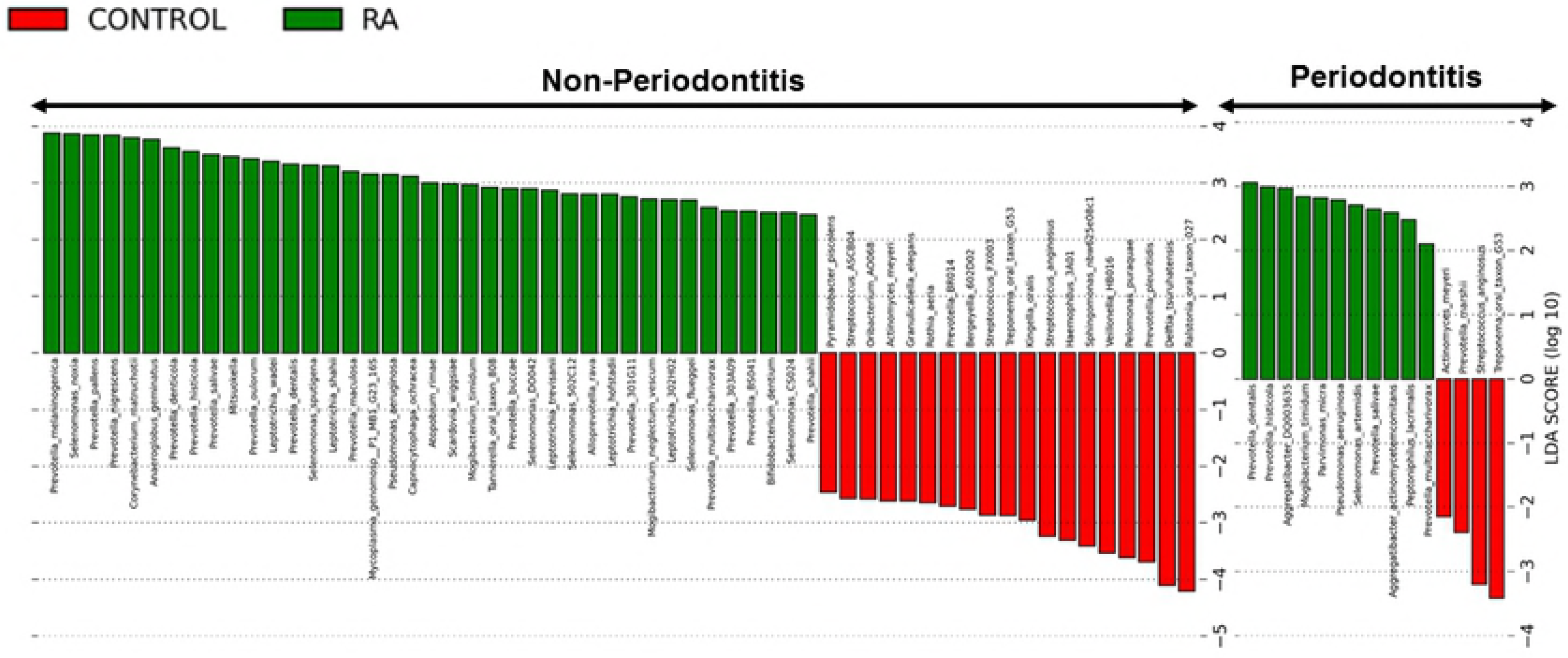
OTUs with different relative abundance based on LEfSe results in Control subjects (Red) and RA patients (Green) without (A) and with periodontitis (B). Bars represent linear discriminant analysis scores (LDA).

We also observe an increased concentration of gram-negative anaerobic species on RA sites compared to control sites with periodontitis (S1 Fig)

### RA parameters and oral bacteria species

Some bacteria species were correlated with RA parameters. In RA subjects with periodontitis, bacteria related to periodontal health, such as *Actinomyces,* were negatively correlated with number of tender joints (rho= −0.36, p<0.05). On the other hand, in the same subjects, the presence of pathogenic species such as *Fretibacterium fastidiosum, Parvimonas micra* and *Anaeroglobus geminatus* were correlated with augmented numbers of swollen (rho= 0.35) and tender joints (rho= 0.30), p<0.05.

### Predicted functional signatures of subgingival microbiota in RA patients

Analysis using PICRUSt revealed that genes involved with energy metabolism, lipopolysaccharide (LPS) biosynthesis, amino acid and carbohydrates metabolism, cell cycle and peptidases were significantly more abundant in the subgingival metagenome of subjects with RA independent of periodontal status. In controls, genes involved with amino acid biosynthesis, and carbohydrate metabolism were overrepresented in the microbiota (S2 Fig).

### Salivary concentration of inflammatory cytokines in RA patients

To investigate whether the above-mentioned dysbiosis in subgingival microbiota could be associated with an altered inflammatory response we measured cytokines in saliva of RA and control subjects (Fig 4). The levels of IL-2, IL-6 and IFN-γ were increased in saliva from RA patients compared to control subjects both without periodontitis (P<0.05). IL-33 and TNF-α were increased in all RA groups independent of periodontal status (P<0.05). IL-17 was increased in RA subjects with periodontitis compared to control subjects (P<0.05, Fig 4).

**Figure 4.**
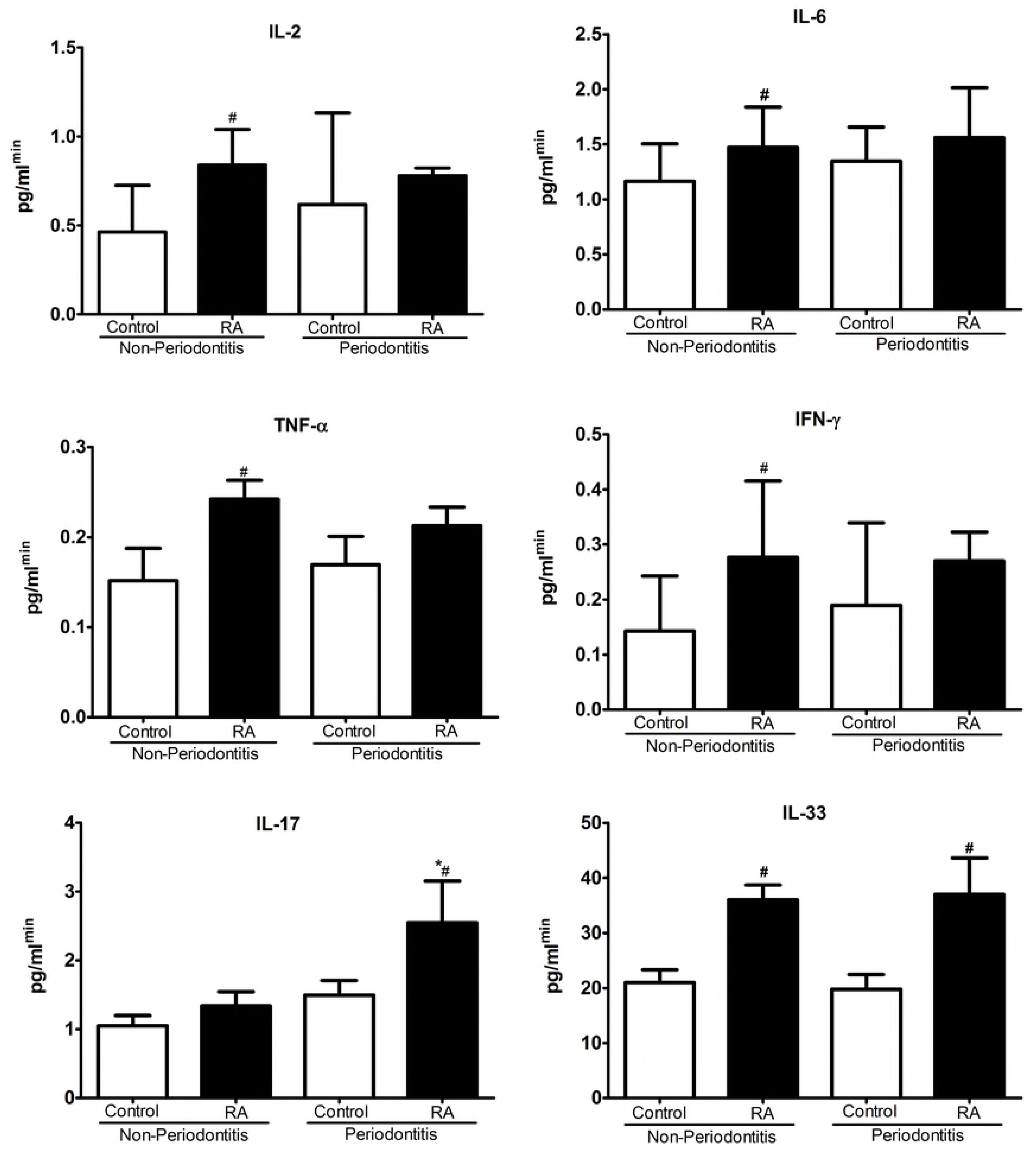
Levels of inflammatory cytokines in saliva. Control subjects and patients with Rheumatoid Arthritis (RA) with and without periodontitis, determined by ELISA and CBA. *Statistically different compared to Non-Periodontitis subjects within the same group. #Statistically different compared to Control. p<0.05, Kruskal-Wallis

The increased levels of the cytokines IL-6, IL-17 and IL-33, positively correlated with periodontal parameters such as probing depth and number of missing teeth as demonstrated in Table 2 (P<0.05). The levels of IL-33 positively correlated with RA parameters such as Rheumatoid factor and c-reactive protein (CRP), while IL-6 was positively correlated with CRP and ESR. Furthermore, the presence of healthy-related species including *Streptococcus, Rothia aeria, Actinomyces* was negatively correlated with cytokines IL-17 and TNF-α. In contrast, the presence of pathogenic species, such as *Selenomas* and *Prevotella,* were correlated with increased levels of inflammatory cytokines (IL-2, IL-6, IL-17, IL-33 and TNF-α) (Table 2).

**Table 2.**
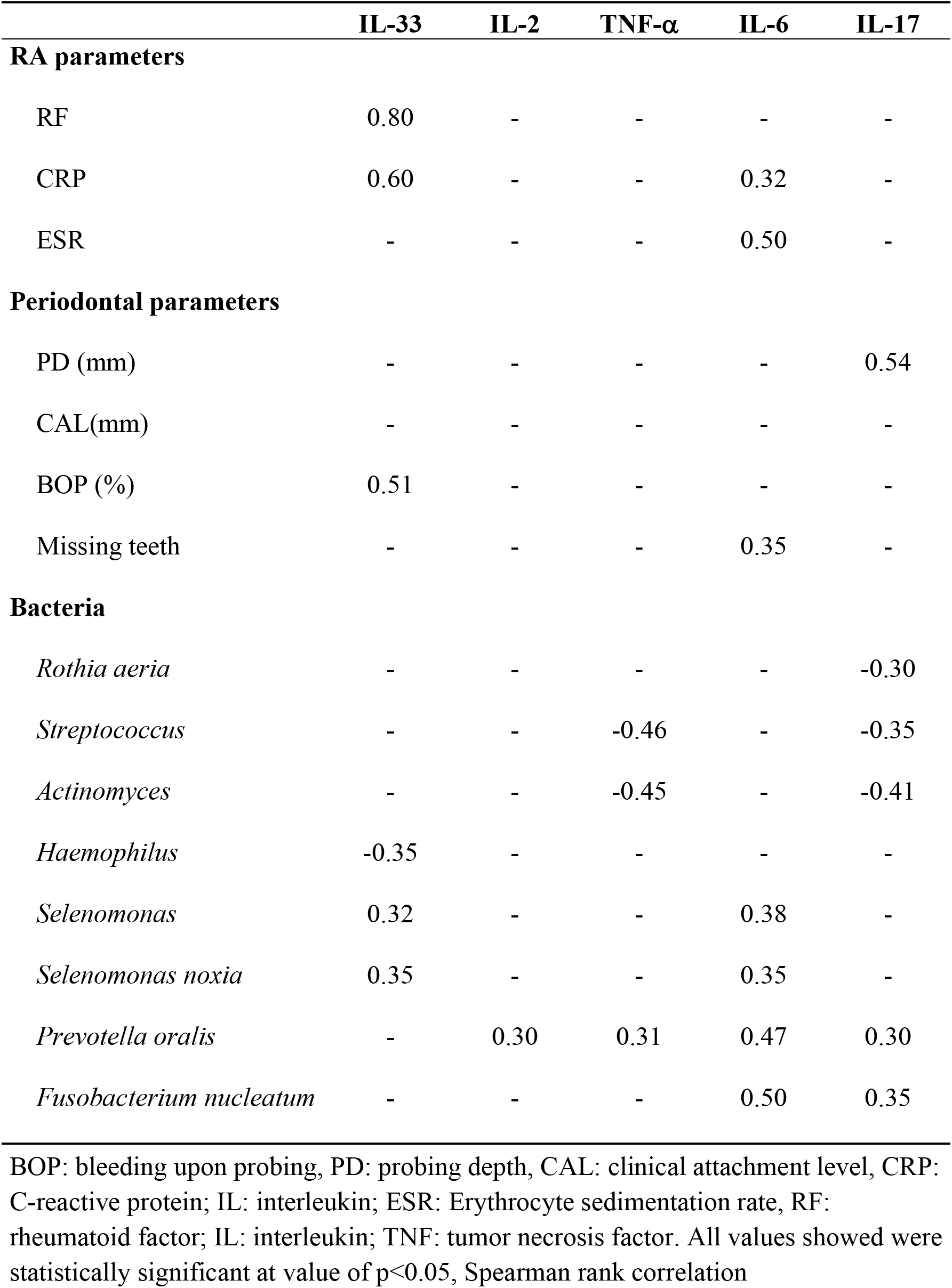
Correlations among inflammatory cytokines in saliva, relative abundance of bacteria, RA and Periodontal parameters in RA patients (rho values)

### Subgingival biofilm of RA patients elicited high IFN-γ production from PBMCs

To evaluate the potential of dysbiotic subgingival biofilm of RA patients to stimulate inflammatory response, we exposed PBMCs from control individuals to inactivated bacterial plaque samples from RA or healthy donors, neither of which had periodontitis. Plaque samples from RA subjects’ stimulated significantly higher production of IFN-γ compared to plaque from control subjects (Supplemental Figure 3).

## Discussion

A relationship between RA and periodontitis has previously been reported, but the impact of RA on the subgingival microbiota linked to periodontal disease has not been thoroughly investigated and mechanisms for the potential impact have not been addressed [23–28]. In the present study, we showed significant differences in the subgingival bacterial community between RA patients and controls. RA patients had a higher bacterial load, a more diverse microbiota and increased abundance of pathogenic species compared to controls, even in periodontally healthy individuals. Accordingly, periodontal destruction (probing depth and clinical attachment loss) was significantly greater in RA subjects.

Microbiota homeostasis can be modulated by the host through several factors, including genetic, environmental and inflammatory [29]. Chronic systemic inflammation, as observed in RA, may affect the levels of inflammatory cytokines in periodontal tissues for instance, increased concentration of cytokines in saliva have been consistently reported for chronic inflammatory diseases such as systemic lupus erythematosus [30] and rheumatoid arthritis [31]. We observed higher levels of IL-2, IFN-γ TNF-α, IL-6, IL-17 and IL-33 in saliva from RA patients compared to control subjects. These cytokines have previously been demonstrated in sites of periodontal inflammation [32–34]. IL-2 and IFN-γ are Th1 cytokines that enhance cell-mediate response [35]. TNF, IL-6 and IL-17 have multiple overlapping functions that contribute to RA and periodontal disease by mediating leukocyte activation and migration, chemokine expression and osteoclast activation [26,36]. IL-33 stimulates Th2 cells to secrete the cytokines IL-4, IL-5 and IL-13. It also induces the release of TNF-α, IL-6 and IL-1 [32]. IFN-γ is primarily produced by Th1 and natural killer (NK) cells, and regulates several aspects of the immune response [46]. IFN-γ, together with cytokines such as IL-17, play a central role in the inflammatory reaction and bone resorption in periodontitis [47]. Here we found high levels of IFN-γ in saliva of RA patients. Moreover, when PBMCs were stimulated with dental plaque from RA subjects we observed a much higher production of IFN-γ in comparison with PBMCs exposed to dental plaques of systemically healthy individuals.

Previous studies have suggested that periodontitis and RA are possibly interdependent with respect to the elevated levels of pro-inflammatory molecules [37]. Inflammatory mediators found in the subgingival microenvironment may change the ecological conditions in favor to the outgrowth of pathogenic bacteria, leading to periodontal destruction [38]. Local inflammation enhanced by systemic disease may change the microbial composition toward one that is adapted to an inflammatory environment and more capable of inducing inflammation, as shown for diabetes [4]. The increased inflammation caused by RA coupled with microbial changes may amplify periodontal inflammation and explain the greater susceptibility to periodontitis that we and others have observed [39,40]. In agreement with this observation, studies in mice showed that RA induces alveolar bone loss which is linked to changes in oral microbiota of these animals [27]. Thus, the results presented here provide further support that similar events occur in humans.

We found that RA was associated with increased microbial load and diversity, which is consistent with previous reports that periodontitis, unlike most polymicrobial infections, is associated with increased bacterial diversity [41]. In addition, we observed that the severity of RA, reflected by the number of tender and swollen joints, was significantly correlated with the presence of pathogenic oral bacteria (i.e. *Fusobacterium nucleatum* and *Treponema socransky*). In agreement with our results Zhang et al. [11] reported that bacteria enriched in RA individuals showed positive correlations with RA parameters. However, this study did not investigate whether differences in the oral microbiota were associated with periodontitis. Another study evaluated only RA patients without periodontal disease [13] and their results are in agreement with our findings as we observed that RA subjects without periodontitis harbored increased bacterial biomass. Interestingly, a general pattern of enrichment of *Prevotella* species in RA patients was noticed. This finding is quite interesting when we look at the metabolic pathways of *Prevotella* species which break down proteins and peptides into amino-acids and degrade them further to produce short-chain fatty acids, ammonia, and sulfur compounds [42]. Our analysis of predicted functions using PICRUSt confirmed an enrichment of several pathways of amino acid metabolism and peptidases in the microbiota from RA subjects. In addition, we observed overexpression of genes linked to bacterial virulence such as LPS and sporulation genes. These metabolites are able to induce tissue inflammation [43] and to promote fibroblasts apoptosis [44], contributing to periodontal destruction. In addition, periodontal inflammation can increase the secretion of gingival crevicular fluid, leading to a protein-rich environment and overgrowth of proteolytic bacteria, such as *Prevotella*, keeping the tissue destruction cycle [42]. In agreement with this, we also found that RA subgingival sites had an increased concentration of gram-negative anaerobic species, that are strongly associated with proteolytic metabolism and consequently, periodontal destruction [45]. In addition, we also found higher levels of *Selenomonas noxia* and *Parvimonas micra* in RA subjects in agreement with previous findings in animal model of RA [27].

Recently our group reported the impact of another systemic autoimmune disease, systemic lupus erythematosus (SLE) on periodontal status and subgingival microbiota [49]. Patients with SLE exhibit a higher prevalence of periodontitis, which occurs at a younger age when compared to healthy individuals. Like RA, SLE caused an increased bacterial load in subgingival sites and induced changes in the microbial composition and diversity that were linked to increased cytokines concentration on saliva (IL-6, IL-17 and IL-33), similar to the changes observed for RA patients. Together these results corroborate the hypothesis that systemic inflammatory conditions lead to dysbiosis of subgingival microbiota and increase the risk of periodontitis.

The limitations of our study rely on its cross-sectional design, with a single time measurement. Therefore, we are unable to answer the question whether changes in the microbiota are a cause and/or effect of the RA. Besides this, the use of different types of medications to treat RA may affect the periodontal inflammation and microbiota. It is also important to mention that the analysis of predicted functions by PICRUST is only descriptive, as the ideal method to assign metabolic functions is direct analysis by RNA sequencing.

In conclusion, our study is the first to demonstrate the influence of RA on subgingival microbiota considering the periodontal status of the subjects. Our findings support the concept that RA is a chronic inflammatory disease that triggers and/or aggravates the imbalance between pathogenic bacteria/health-related bacteria in subgingival biofilm thus increasing susceptibility to periodontal diseases.

## Methods

### Subjects

During one year we evaluated patients from the Rheumatology Outpatient Clinic of Clinics Hospital of Federal University of Minas Gerais (UFMG), Belo Horizonte, Brazil that were diagnosed with RA based on the 2010 American College of Rheumatology and EULAR classification criteria [14]. Two hundred thirty-nine patients agreed to participate, however, the study group consisted of forty-two patients based on the following inclusion criteria: no other rheumatic disease, no treatment for periodontal disease within the last 6 months, no use of orthodontic appliances, no use of antibiotics within the last 3 months, no pregnancy or lactation and the presence of at least 8 teeth. The control group consisted of 47 subjects without RA or other rheumatic diseases, that were age and gender matched with the RA group, randomly assigned from a population with demographic, social, and educational backgrounds similar to RA patients. Their medical history was obtained from an interview. Patients’ medical history and medications were determined by review of medical charts. Each patient had laboratory assessments of blood levels of IgM rheumatoid factor (RF), C-reactive protein (CRP), anti–citrullinated protein antibody (ACPA) and erythrocyte sedimentation rate (ESR). Periodontal status was assessed by two calibrated examiners (JDC and SMM) and the following parameters were recorded: plaque index, probing depth, clinical attachment level and bleeding on probing. Periodontitis was defined as the presence of two or more interproximal sites with probing depth ≥ 4 mm or one site with probing depth ≥ 5 mm [15].

## Ethics Statement

The subjects gave written informed consent, and the study protocol was approved by the Federal University of Minas Gerais Ethics Committee (CAAE: 03128012.0.0000.5149/2012). All patient data and subgingival samples were anonymized. For sampling of PBMCs from the blood of healthy subjects they also gave written informed consent and their data were anonymized.

### Subgingival samples collection

Subgingival samples were collected using endodontic paper points (ISO40) (Tanariman, Manacaparu, AM, Brazil) that were inserted in 5 sites with deepest probing depth for one minute. After removal, the material was pooled together and stored in a sterile tube containing 500 µL of sterile distilled water and centrifuged at 3,000 g for 5 minutes. The paper points were discharged, and the pellet was kept at −80°C until DNA extraction.

### Saliva collection

Saliva was collected by continuous drooling into a sterile 50 mL tube for 5 minutes. The salivary flow was measured in milliliters per minute (ml/min). The saliva samples were diluted (1:1) in a phosphate-buffered saline (PBS) solution containing protease inhibitors and subsequently frozen at −80°C until analysis.

### Cytokines measurement

Concentrations of interleukin-2 (IL-2), IL-6, IL-17, tumor necrosis factor-α (TNF-α and interferon-γ (IFN-γ) in saliva were determined using a Cytokine Bead Array (CBA) Human Th1/Th2/Th17 Kit (BD Biosciences, San Diego, CA) and analyzed on a BD FACS Calibur flow cytometer (BD Biosciences). The concentration of IL-33 was measured by enzyme-linked immunosorbent assay (ELISA) (R&D Systems, Minneapolis, MN, USA). Assays were performed according to the manufacturer’s instructions. The results were expressed as picograms of cytokine (pg/ml) adjusted according to salivary flow.

### DNA Extraction and Sequencing

DNA was extracted from plaque samples using the Quick-g DNA MicroPrep kit (Zymo Research, Irvine, CA, USA) and 50 µL (10 mg/ml) of lysozyme per sample to enhance bacterial lysis as described [16]. The quantity and quality of DNA was measured spectrophotometrically (Tecan, Männedorf, Switzerland). The primers 515F (5’ -GTGCCAGCMGCCGCGGTAA-3’) and 806R (5’ -GGACTACHVGGGTWTCTAAT- 3’) which target the hypervariable V4 region of the 16S rRNA gene were used for amplification [17]. After, agarose gel electrophoresis was performed to check size integrity. All amplicons were subjected to Illumina MiSeq Platform at the Next-Generation Sequencing Core of University of Pennsylvania and sequenced together at the same run. All Illumina sequence data were submitted to the NCBI Sequence Read Archive (SRA) under BioProject accession number PRJNA325500.

### Microbiota Analysis

The raw reads were trimmed to remove regions with a low Phred score. Trimmomatic [18] was the tool used with the TRAILING:5 and SLIDINGWINDOW:4:15 parameters. The trimmed reads were merged using FLASH [19] tool requiring 30 reads overlap. The assembled amplicons were mapped to the CORE database using the Qiime’s pick_closed_reference_otus and 97% identity threshold. The representative set of sequences had their taxonomic classification using the same database and the Qiime’s assign_taxonomy script. The alpha diversity indexes were assessed using the Vegan R package [20]. Beta diversity was calculated using Unique Fraction metric (UNIFRAC) unweighted. Quantification of total bacterial load was determined by real-time PCR using universal primers for 16S rRNA gene. (F: AGAGTTTGATCCTGGCTCAG; R: ACGGCTACCTTGTTACGACTT) (IDT, Coralville, Iowa, USA) based on a a standard curve prepared using DNA extracted from a known number of *Porphyromonas gingivalis* (colony forming units) separated with flow cytometry and amplified with the same qPCR protocol. Samples were assayed in duplicate in a 25 μl reaction mixture containing 2.5 μl of template DNA, 2.5 μl of 10x TaqMan Universal PCR Master Mix, 1.5 pl of MgCl_2_,1 pl dNTP, 12.5 pmol of forward primer and reverse primer. The standard curve was used to derive the Cq (quantification cycle value) vs log CFUs linear equation. The cycling conditions used were as follows: 95 °C for 10 min, followed by 40 cycles at 95 °C for 15 s and 60 °C for 1 min each.

To provide an inference of the functional profile of the microbial community based on 16S rRNA gene sequence results we assigned taxonomy to the representative set of sequences using the GreenGenes 13.5 database. This classification was utilized by PICRUSt (phylogenetic investigation of communities by reconstruction of unobserved states) to identify and quantify the Pathways and KEGG Orthology Groups.

### Blood collection and in vitro induction of PBMCs with oral biofilm

Blood was obtained from 5 systemically healthy individuals that did not have periodontitis or current use of immunosuppressive or anti-inflammatory drugs. Peripheral blood mononuclear cells (PBMCs) isolated by Ficoll–Paque gradient (Amersham Biosciences, Uppsala, Sweden) for 40 minutes at 20ºC. PBMCs were washed twice with PBS and counted in a hemocytometer chamber. The PBMCs of each patient (106 cells/ml) were incubated in a complete RPMI medium of 2 mM L-glutamine, 5% normal human serum, 100 l g/mL streptomycin, and 100 UI/mL penicillin G potassium with subgingival plaque in 96-well plates. For these assays total subgingival plaque was removed from a given site, placed in 100 ul of TE buffer, homogenized by vortexing and heat/freeze inactivated. PBMCs were exposed to a standardized amount of bacteria (1×10^8^ CFU) for 24 h and then supernatants were processed to evaluate cytokine concentration.

### Statistics

After evaluating the normality of distribution by Kolmogorov-Smirnov tests, clinical, demographic, alpha diversity, cytokine levels and bacterial load data were compared using One Way ANOVA or Kruskal-Wallis test. Correlations between relative abundance of taxa and clinical parameters of periodontal disease and RA were calculated using Spearman correlation coefficients (SPSS software, version 20). PERMANOVA was performed to compare beta diversity (QIIME software, version 1.9). In case of multiple comparisons, the p-value was corrected using Bonferroni correction. The predicted functional groups, and Operational Taxonomic Units (OTU), were compared among RA and control subjects and tested for statistical significance using DESeq2 [21] (R statistical software, version 3.5) and LEFSE [22], respectively. P values <0.05 were statistically significant.

## Data reporting

All Illumina sequence data from this study were submitted to the NCBI Sequence Read Archive (SRA) under BioProject accession number PRJNA325500.

## Acknowledgements

We thank Professor Dr. Paulo Eduardo Alencar de Souza for his contribution with in vitro experiment.

## Supporting information

**S1 Fig. – Relative abundance (%) of microbiota composition.** Gram status and oxygen metabolism of subgingival microbiota in RA patients and Control subjects, without and with periodontitis.

**S2 Fig – Differentially abundant gene functions in subgingival microbiota.** Control and RA subjects without (A) and with periodontitis (B). Functional categories of genes of the subgingival metagenome were predicted by using PICRUSt, and differentially abundant functions were then identified by using linear discriminant analysis (LDA) coupled with effect size measurements (LEfSe).

**S3 Fig – Immunostimulatory potential of dental plaque from RA patients.** Exposure of human PBMCs to RA microbial plaque. Levels of IFN-γ was determined by ELISA. statistically different compared to Control. p<0.05, Student t-test.

